# Visual Processing During the Interictal Period Between Migraines: A Meta-Analysis

**DOI:** 10.1101/2021.09.13.459979

**Authors:** Timucin Sezai, Vinh Nguyen, Nina Riddell, Melanie J. Murphy, Sheila G. Crewther

## Abstract

Migraine is a poorly understood neurological disorder and a leading cause of disability in young adults. Migraines are characterized by severe pulsating unilateral headache and visual symptoms. Whether visual function is also impaired in the interictal period between migraines remains controversial. Thus, this meta-analysis investigated the evidence for altered visual function as measured electrophysiologically via pattern-reversal visual evoked potential (VEP) amplitudes and habituation in adult migraineurs with or without visual aura and control in the interictal period. Twenty-three studies were selected for random effects meta-analysis, demonstrating slightly diminished VEP amplitudes and substantially reduced habituation in the early P100 component in migraineurs without aura and with aura compared to controls. No differences were found between migraineurs with and without aura. Although heterogeneity between studies and insufficient published data for VEP latencies and the earlier N75 VEP component data was observed and require further testing, P100 anomalies may indicate abnormal functioning of the fast-conducting magnocellular visual pathway, in episodic migraineurs.

## 1. Introduction

Migraine is an underdiagnosed and inadequately managed condition (Lipton et al., 2013) characterised by periodic severe recurring headaches and neurological symptoms that cause extended periods of disability (International Headache Society [IHS], 2018). Migraine is the second most prevalent neurological condition across the adult lifespan, the most prevalent neurological condition in young working age adults (Stewart et al., 2008), and the sixth leading cause of disability worldwide (Vos et al., 2017). A typical migraine involves pulsating headaches with moderate to severe unilateral pain, and can be accompanied by nausea, vomiting, and/or sensitivity to light (photophobia) and sound (phonophobia; Goadsby et al., 2017; IHS, 2018). The International Classification of Headache Disorders 3rd edition (ICHD-3) categorises migraine as two primary disorders: migraine without aura (MO) and migraine with aura (MA), where aura is defined as sensory symptoms that are predominantly visual, arising shortly before or during a headache (IHS, 2018). The MA subtype is primarily characterised by more extreme visual symptoms, often including perception of zigzag patterns and/or scintillating scotoma (holes in vision) that gradually spread across the visual field (IHS, 2018). Thus, the present study aimed to meta-analyse the electrophysiological literature associated with visual function in migraineurs to determine whether visual function is only perturbed during migraine, or more permanently altered during the interictal period (i.e., between migraine events) in episodic migraineurs.

Vision is the primary spatial and temporal information processing modality of the primate brain (Felleman & Van Essen, 1991; Livingstone & Hubel, 1987) driving attention, cognition, and goal-directed actions in humans (Corbetta & Shulman, 2002; Crewther et al., 2012; Laycock et al., 2007). Visual processing and eye movements also occupy large cortical (Felleman & Van Essen, 1991) and subcortical volume (Wurtz & Goldberg, 1971) and require the greatest proportion of the brain’s metabolic resources (Wong-Riley, 2010). Visual symptoms are among the core diagnostic criteria of migraine, yet the role of interictal visual abnormalities in migraine pathophysiology remains unclear. Two interconnected theories that describe possible visual processing anomalies in migraineurs during the interictal period relate to visual system *(i) excitability* and *(ii) habituation* to repetitive stimulation (Magis et al., 2013).

Although heightened sensitivity to visual stimulation is predominantly experienced during a migraine episode, this has also been observed in some migraineurs during the interictal period (Peroutka, 2014; Shepherd, 2019), leading to the hypothesis that the visual system in migraineurs is abnormally sensitive or hyperexcitable to visual stimulation both during and outside of a migraine event (Aurora & Wilkinson, 2007). Vecchia and Pietrobon (2012) have further suggested that hyperexcitability and imbalances in visual cortex excitatory and inhibitory mechanisms contribute to susceptibility to cortical spreading depression (CSD) in MA. This hypothesis is supported by meta-analyses of studies examining cortical arousal levels in interictal migraineurs and non-migraine controls using transcranial magnetic stimulation (TMS), finding primary visual cortex (V1) hyperexcitability in those with MA, but not MO (Brigo et al., 2013). While meta-analysis is available summarising TMS in migraine populations, no current attempts have been made to systematically summarise other electrophysiological methods, such as visually evoked potentials (VEPs) recorded using electroencephalography (EEG). Given the frequency of visual anomalies in migraine, electrophysiological analysis of visual system functioning during the interictal period between migraines is important for understanding migraine pathophysiology and its effects on baseline function and cortical excitability. Moreover, the high temporal and reasonable spatial resolution of the VEP technique allows objective quantification of the effects of cortical excitability in susceptibility to migraine (Magis et al., 2013).

There is little consistency across studies recording VEPs in migraineurs during the interictal period, particularly for studies analysing VEP component amplitudes and latencies (Ambrosini et al., 2003). To address this, Odom et al. (2016) have suggested that between-participant variability in VEPs may be minimised by using a black-and-white checkerboard pattern-reversal stimulus that inverses at a transient rate (approximately three or less reversals per second) whilst recording responses from an occipital electrode above V1. The majority of studies available using a checkerboard pattern-reversal VEP methodology in migraine populations have also reported the P100 and N135 time point components (Coppola et al., 2019) based on Klistorner et al. (1997) use of temporal analysis to dissociate separate magnocellular (P100-N115) and parvocellular (N100-P120-N160) contributions to the human multifocal flash VEP (mfVEP). Existing VEP studies of interictal migraineurs have identified greater responses (hyperexcitability), diminished responses (hypo-excitability) and even normal responses when compared to non-migraineurs (Magis et al., 2016; Tolner et al., 2019). Furthermore, the role of methodological and confounding factors on electrophysiological response in migraineurs, such as time in the migraine cycle (Coppola & Schoenen, 2012; Schoenen, 2011; Stankewitz & May, 2009), preventative medications (Magis et al., 2013), participant age (Brown et al., 2019) and VEP stimulus size and speed (Odom et al., 2016), is poorly understood. Factors such as VEP stimuli and time of assessment with reference to the migraine cycle and time point during the interictal period varies significantly across previous studies (Ambrosini et al., 2003; Magis et al., 2007). A review by Magis et al. (2016) discussed findings across all visual electrophysiology methods in migraine post-2013, but only reported significant results and did not describe how studies were selected for review. Finally, a very recent review by de Tommaso (2019) summarised spectral analysis EEG and steady-state VEPs in migraine, but not transient VEPs as a means of understanding the impact of migraine on speed of visual processing. Thus, the literature remains highly controversial with conflicting evidence and inconsistent methodologies used to analyse VEPs in interictal migraineurs.

Electrophysiological function in migraineurs has also been associated with abnormal responses to prolonged stimulation and habituation mechanisms across visual, auditory, somatosensory, and nociceptive evoked potentials (Brighina et al., 2009; Magis et al., 2013). Habituation in VEP studies refers to a progressive decline in response amplitude across prolonged stimulation (Omland et al., 2011), and is often interpreted as the reduction in attention required to respond to a non-changing stimulus (McDiarmid et al., 2017). Abnormal habituation in migraine has been linked to abnormal cortical excitability and vulnerability to sensory overload, and may be a consequence of repeated exposure to pain/stress activation caused by migraine symptoms (Stankewitz & May, 2009). Repeated cortical inflammation and pain activation associated with frequent migraines may alter the cortical resting state, causing abnormal electrophysiological responses in interictal migraineurs (Kowacs et al., 2015). If so, more frequent migraines and greater time between onset of regular migraines could moderate VEP habituation, amplitudes, and latencies (Nguyen et al., 2012). However, habituation deficits are not consistently identified in interictal migraineurs, possibly due to participant or methodological differences across studies (de Tommaso et al., 2014). Potentially the most important factor affecting VEP habituation patterns is the stage of the migraine cycle at which recordings are obtained (Coppola & Schoenen, 2012). As such, inconsistent findings across studies raises questions as to whether reduced habituation is a useful measure to understand the cortical and visual contributions to migraine aetiology, vulnerability, or treatments during the quiescent interictal period.

Thus, the present study aimed to examine visual processing in migraineurs during the interictal period by conducting a meta-analysis on case-control studies that have compared VEP responsivity and habituation in adult migraineurs and non-migraine controls. The foci of the meta-analysis were the traditional P100 and N135 timepoints measured using checkerboard pattern-reversal VEP stimuli, which are the most commonly available results. It was hypothesised that meta-analysis would demonstrate changes in early VEP amplitude and decreased habituation (i.e., smaller response decrement to prolonged stimulation) in individuals diagnosed with migraine compared to non-migraine controls. Our secondary aims were to examine differences between MA and MO subtypes and, where possible, explore potential confounds of VEPs in migraine including: Age, migraine frequency, years suffering migraines, and VEP stimulus check size and reversal rate.

## 2. Method

### 2.1. Search Strategy

This study was pre-registered with Open Science Framework on May 25th, 2020 (registration doi: 10.17605/OSF.IO/VHWEP). Searches of MEDLINE, Embase, PubMed and Web of Science, PsycINFO and CINAHL databases were conducted for peer-reviewed studies written in the English language with no date restrictions published up to June 29th, 2021. Title and abstract search terms were: 1) “migraine” OR “migraine aura” OR “migraine headache” OR “headache disorder”; and 2) “VEP” OR “visual* evoked potential*” OR “VER” OR “visual* evoked response*” OR “functional vision”; and 3) “EEG” OR “electroencephalogra*”, incorporating Medical Subject Headings (MeSH) where possible. Reference lists of extracted studies were also manually searched. Results were imported into Covidence Systematic Review Software (“Covidence”; Veritas Health Innovation, 2019).

### 2.2. Study Selection Criteria

Primary inclusion criteria were case-control studies with a between-groups design comprising adults (18-60 years of age) with migraine and non-migraine controls being compared in VEPs recorded with EEG. Studies primarily used the ICHD (IHS, 2018) to diagnose migraineurs with “migraine without aura” (MO) or “migraine with aura” (MA). Non-migraine headache disorders (e.g. tension-type headache) were excluded. Included studies required a healthy control group without migraine or headache history and screening participants for neurological disorders with abnormal electrophysiological responses (e.g. epilepsy) that could confound VEPs (Vialatte et al., 2010). Adult migraineurs aged 18-60 were selected to reduce age-related confounds on VEPs, as previous research suggests VEP amplitudes and latencies decrease across the lifespan (Brown et al., 2019). Age ranges were requested from authors if missing from articles and studies were excluded if one standard deviation of the mean age lay outside 18-60. Included studies explicitly tested interictal migraineurs free from migraine symptoms for at least 48 hours before and after testing. Studies where migraineurs used preventative medication during testing were excluded as this can impact VEPs (Coppola et al., 2009).

Methodological inclusion criteria were that VEPs were recorded using EEG according to international standards (Odom et al., 2016). The International Society for Clinical Electrophysiology on Vision states that VEPs require an active electrode on the occipital scalp (“Oz”) for recording the visual cortex in line with the 10-20 international system (Odom et al., 2016). Inclusion was limited to studies using checkerboard pattern-reversal VEP stimuli due to insufficient studies available using pattern onset/offset or flash stimulation or multifocal stimuli. Only transient VEPs were included within the scope of this meta-analysis. Steady-state VEPs and EEG spectral analysis were excluded but may answer future questions related to underlying brain activity in migraine (Vialatte et al., 2010). Thus, included studies assessed the main components of the transient pattern-reversal VEP waveform with amplitude, latency, and habituation (change in baseline wave amplitudes across testing) as outcome measures. Baseline data was obtained where studies recorded VEPs on multiple occasions and studies were excluded if baseline values were unobtainable. See Figure 2 PRISMA flow chart for exclusion criteria and frequency.

**Figure 1.**
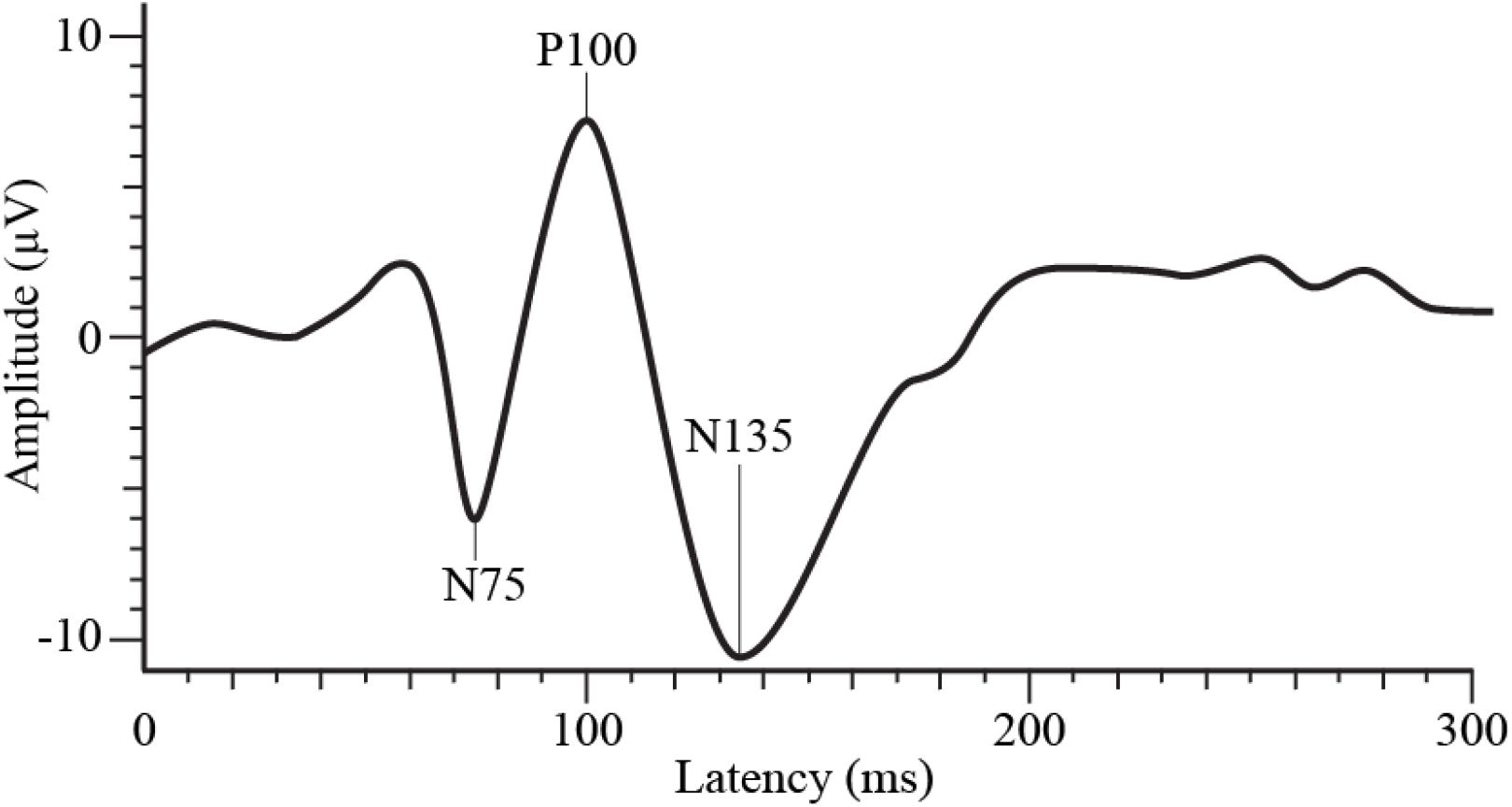
Typical Transient Pattern-Reversal Visual Evoked Potential Waveform *Note*. μV = microvolts, ms = milliseconds. Adapted from “ISCEV standard for clinical visual evoked potentials: (2016 update)” by J. V. Odom et al., 2016, *Documenta Ophthalmologica, 133*(1), p. 7 (https://doi.org/10.1007/s10633-016-9553-y).

**Figure 2.**
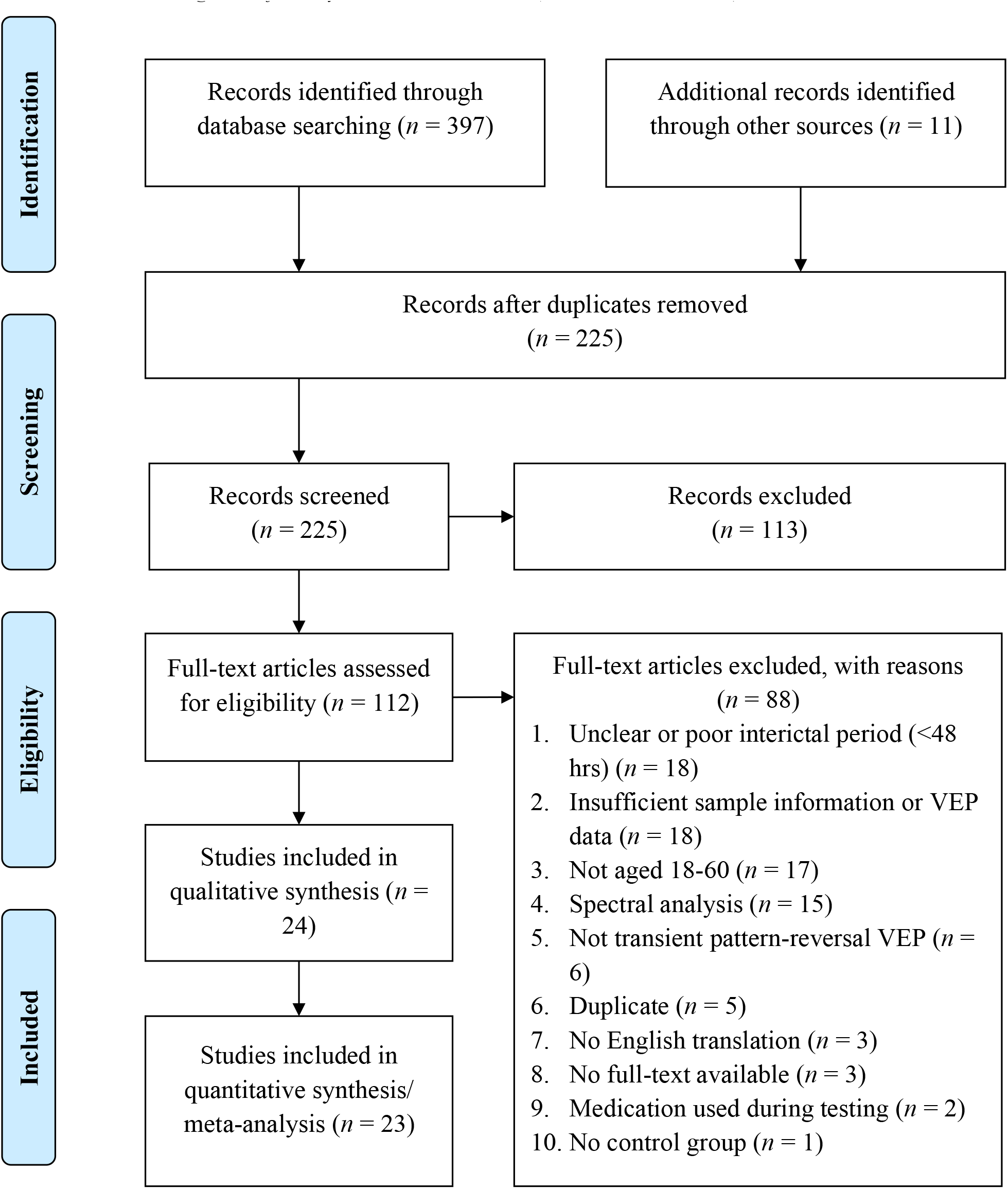
PRISMA Flow Diagram of Study Selection Process (Moher et al., 2009) *Note. n* = number of studies.

### 2.3. Data Collection/Extraction

After TS, NR and VN independently screened abstracts, two independent reviewers (TS and VN) extracted data from articles and resolved discrepancies with SC to create a single Microsoft Excel extraction spreadsheet for the variables summarised in Table 1.

**Table 1.** Data Extracted from Studies Included in the Meta-Analysis

| Category | Variables extracted |
| --- | --- |
| Demographics | Sample size ($N$), age ($M$, $SD$, range), and sex ($n$ of males, females) were extracted for each migraine sample and controls. |
| Migraine sample characteristics | Diagnostic method (e.g. ICHD-3), migraines per month ($M$, $SD$), years with migraines ($M$, $SD$), interictal timeframe (minimum number of hours or days participants were migraine-free before and after testing). |
| Stimulus details | Eye tested (monocular, binocular), spatial frequency (check size; ° [minutes of arc]), temporal frequency (reversals per second [rps]), number of stimulus repetitions (trials) per blocks (blocks x trials). |
| VEP outcome measures for each component ($M$, $SD$) | Amplitude (microvolts; $\mu V$), and habituation (regression slope or percentage change in amplitude from the first to final blocks of presentation; positive value = amplitude increment; negative value = amplitude decrement). |

*Note.* Due to insufficient data in most studies, latencies are not reported.

Where possible, data was extracted separately for MO and MA. When type of migraine was not reported, data was extracted under a Migraine Grouped (MG) variable. Demographics were extracted for each sample to check eligibility and to ensure subsamples were comparable. Migraine characteristics and stimulus details were extracted as potential moderators. Outcome variables extracted were VEP amplitudes and habituation for the P100 and N135 components. As depicted in Figure 1 below, P100 was defined as the positive peak occurring approximately 100ms post-stimulus onset and N135 was defined as the second negative peak occurring between 120-150ms post-onset (Odom et al., 2016).

Depending on data available, VEP amplitudes were extracted as either first block average or grand average across all blocks of testing. Habituation was calculated as either percentage amplitude change between the first and last blocks of testing, or the regression slope of amplitude changes across all blocks of testing (Omland et al., 2011). Where both percentage and slope were provided, percentage change was extracted as this was the most frequent habituation index reported. Authors were emailed requesting VEP data that was missing, unclear, or only displayed in graph form. Where such data could not be provided, Engauge Digitizer (version: 12.1; Mitchell et al., 2019) was used to digitally estimate means and standard deviations from graphs, given that digital extraction has been shown to be more reliable than manual estimation (Jelicic et al., 2016).

### 2.4. Risk of Bias

Risk of bias assessment was conducted in Covidence using the National Heart, Lung and Blood Institute (NHLBI) Quality Assessment of Case-Control Studies tool, which was developed for assessing specific risks of bias associated with drawing conclusions on clinical populations compared to non-clinical controls (National Heart, Lung, and Blood Institute [NHLBI], 2014). The complete risk assessment of the selected studies is provided in Supplementary Table 1.

### 2.5. Data Analysis

Data was analysed using JASP (version 0.13.1; JASP Team, 2020). Meta-analysis was used to assess differences in P100 and N135 amplitudes and habituation between groups. Four subgroups were compared: 1) All migraineurs (MG) versus healthy controls (HC); 2) Migraineurs without aura (MO) versus HC; 3) Migraineurs with aura (MA) versus HC; and 4) MO versus MA. Hedges’ *g* effect sizes were calculated for each two-group comparison within a study (Hedges, 1981). Where means and/or standard deviations were unavailable, other statistics (e.g. *t* statistic, confidence intervals) were converted to calculate effect sizes according to the Cochrane Handbook (Higgins et al., 2020). One study contributed two sets of effect sizes for different sized stimuli (Omland et al., 2013), as such the larger check size was included in P100 analysis and the smaller in N135 based on previous literature (Klistorner et al., 1997).

A random effects meta-analysis was conducted using the restricted maximum likelihood method (Kalaian & Raudenbush, 1996; Kontopantelis & Reeves, 2012). Meta-effects with 95% confidence intervals were calculated for each subgroup and visualised using forest plots. Heterogeneity between studies was assessed using *I*^2^ and was interpreted according to effect sizes and evidence for heterogeneity (Higgins et al., 2020). Where heterogeneity was significant (*p* < .05), meta-regression was conducted using migraine characteristics or stimulus details as moderator variables depending on available data. Significant meta-analyses were screened for publication bias using Rosenthal’s fail-safe *N* (Rosenberg, 2005; Rosenthal, 1979) and assessment of funnel plot asymmetry using rank correlation and Egger’s regression significance tests (*p* < .05). Funnel plots and asymmetry tests are provided for all analyses in Supplementary Figures 6-13 and were only interpreted and reported in meta-analyses when ten or more studies existed (Sterne et al., 2011).

## 3. Results

### 3.1. Study Selection

Initial database searching identified 225 studies, of these 112 were retained for full-text screening, after which 88 were excluded from the analysis for the following reasons: 18 studies tested migraineurs within 48 hours of a migraine or did not clearly report testing interictal migraineurs, 18 lacked sufficient sample information to assess eligibility (e.g. failure to disclose absence of epilepsy) or lacked usable VEP data, 17 included participants aged outside 18-60, 15 performed EEG spectral analysis not applicable to the current study, 6 did not use transient pattern-reversal VEPs (e.g. steady-state or flash, 5were duplicates, 3 lacked English translation, 3 lacked available full-text, 2 reported use of migraine preventatives during testing, and 1 lacked non-migraine controls. Eleven authors were emailed requesting additional data across 22 potentially eligible studies. Four provided data, one was unable to locate the data, two authors could not be contacted via email, and four did not respond. Risk assessment was performed on 24 studies and one study was excluded with high risk of bias. The final sample comprised 23 studies containing data for one or more analyses. Figure 2 presents a Preferred Reporting Items for Systematic Reviews and Meta-Analyses (PRISMA) flow diagram (Moher et al., 2009) depicting the number of studies obtained from searches and screened for inclusion.

### 3.2. Risk of Bias Assessment

Risk assessment was conducted on 24 studies using the NHLBI Quality Assessment of Case-Control Studies tool (NHLBI, 2014) and 15 studies were rated good quality with low risk of bias. Six were fair quality due to unclear interictal period description (Áfra et al., 1998), unclear or inconsistent method of migraine diagnosis (Ambrosini et al., 2016a; Ambrosini et al., 2016b), vague description of control group (Coppola et al., 2010b; Ince et al., 2017) or unclear exclusion of comorbid disorders (Ozkul & Bozlar, 2002). Two studies were rated poor quality due to incomplete description of the control group (Judit et al., 2000) and ambiguously described interictal period (Logi et al., 2001), but were not deemed to be outside the parameters of the inclusion criteria. One study was deemed high risk of bias and excluded due to merging P100 and N135 data (Sand & Vingen, 2000), which did not fall under any particular risk assessment category. Following risk assessment, 23 studies were included in the meta-analysis. See Supplementary Table 2 for detailed results of the risk of bias assessment.

### 3.3. Study Characteristics

**Table 2.** Descriptive Statistics for Sample Sizes Across Subgroups for the Included Studies

| Sample | Total *N* | Range | Median *N* | *M* | *SD* |
| --- | --- | --- | --- | --- | --- |
| MG | 897 | 8-280 | 42 | 56 | 62 |
| MO | 359 | 13-44 | 21 | 22 | 11 |
| MA | 170 | 8-35 | 17 | 18 | 8 |
| HC | 766 | 8-240 | 24 | 33 | 46 |

*Note.* *N* = number of participants, Range = minimum-maximum, MG = all migraineurs, MO = migraineurs without aura, MA = migraineurs with aura, HC = healthy controls.

Meta-analyses included 23 studies published between the years 1998 and 2018, providing 108 effect sizes across P100 and N135 amplitudes and habituation. Supplementary Table 3 lists the studies that contributed effect sizes for each meta-analysis. Of these studies, seven diagnosed migraineurs using the ICHD-1 (30.43%), nine used the ICHD-2 (39.13%), one used both ICHD-1 and ICHD-2 (4.35%), four used the ICHD-3 beta (17.39%) and two did not report a specific diagnostic tool (8.70%). No study contained chronic migraineurs. Thirteen studies exclusively recruited migraineurs from hospitals (56.52%) while the remaining studies used combinations of hospitals, universities, advertisements, and/or participant databases. Age and sex differences between samples were statistically controlled for in only one study, therefore neither could not be used as moderators. Fifteen studies reported frequency of migraines per month (65.22%) and 15 reported years suffering migraines (65.22%) in migraine participants, thereby leaving insufficient studies to analyse these moderators. Eighteen studies tested migraineurs with an interictal interval of 72 hours or greater before and after testing (78.26%), three used a 48 hours interictal period (13.04%), and two were ambiguous (8.70%).

Monocular stimulation was used in 22 studies (95.65%) and binocular stimulation in one study (4.35%). Only 12 monocular studies reported which eye was tested (52.17%), although none reported whether eyes stimulated were selected in relation to the brain hemisphere associated with migraine symptoms. Seventeen studies incorporated small stimuli with check sizes ranging 0.13-0.27° (73.91%), four used large stimuli ranging 0.80-1.13° (17.39%), one used two different sized stimuli (4.35%), and one did not report check size (4.35%). Seventeen studies used reversal rates of 3.1rps (73.91%), three used 3rps (13.04%) and three used 2rps (13.04%), therefore reversal rate was not used as a moderator due to insufficient variability. Supplementary Table 4 provides the study characteristics extracted from articles and figures or supplied by authors. Supplementary Table 5 details the individual results from each study.

### 3.4. Meta-Analysis of Visual Evoked Potentials in Migraine

#### 3.4.1. Meta-Analyses for VEP Amplitudes in Migraine

Random effects meta-analysis using the restricted maximum likelihood method of studies comparing all migraineurs (MG) to healthy controls (HC) in P100 and N135 VEP amplitudes showed no significant differences in P100 amplitude between MG and HC across 17 studies, *g* = −0.15, 95% C.I. [−0.30, 0.01], *p* = .06, *I*^2^ = 26.69%, *p* = .12. No significant differences were observed in N135 amplitude between MG and HC across six studies, *g* = −0.32, 95% C.I. [−0.83, 0.19], *p* = .22, *I*^2^ = 81.50%, *p* < .001. Although there was substantial variance between studies, there were too few studies reporting all potential moderators to perform moderator analysis. Figure 3 provides forest plots illustrating the results of differences between MG and HC in VEP amplitudes for both waveform components.

**Figure 3.**
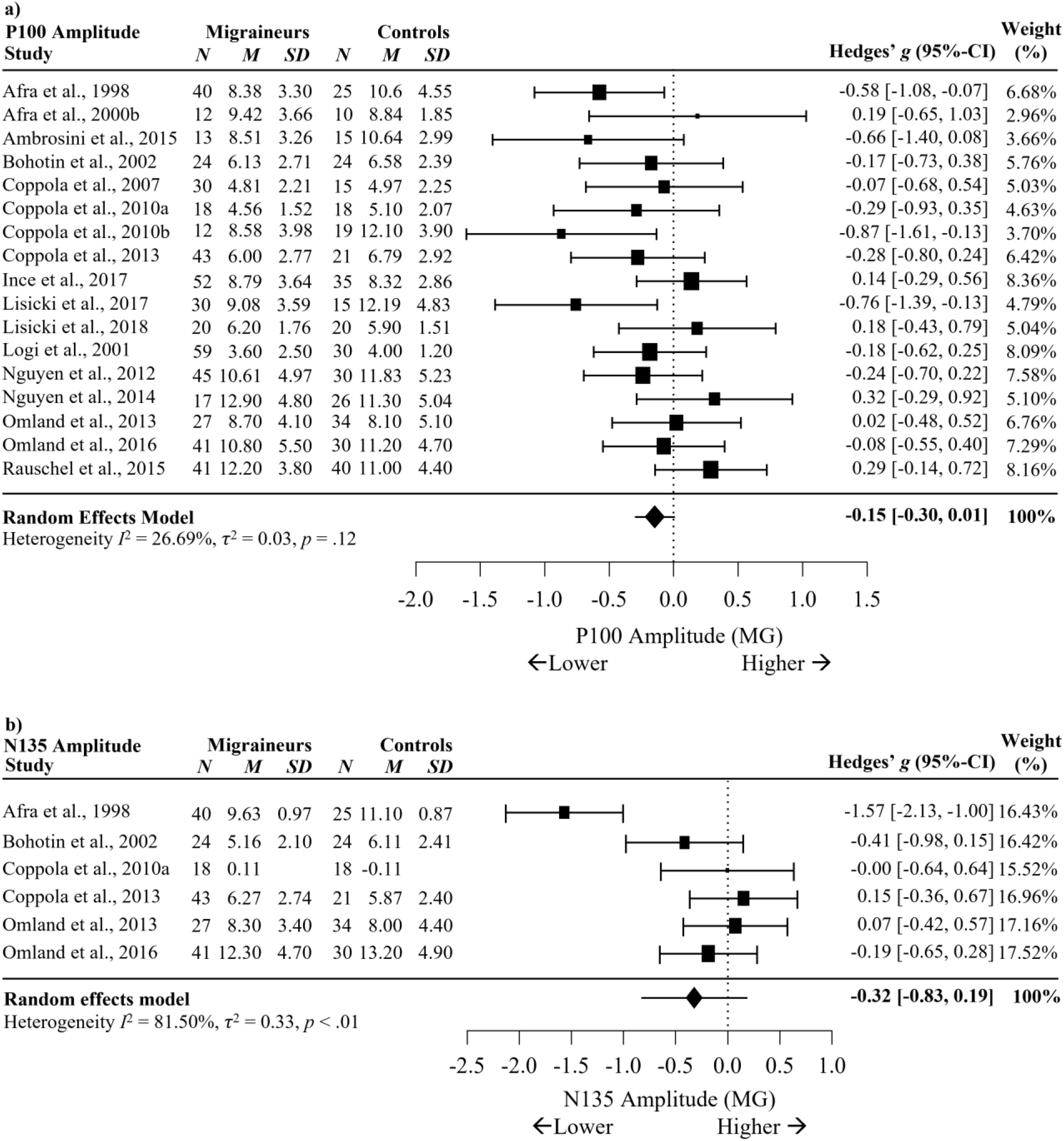
Forest Plots Comparing a) P100 and b) N135 Amplitude during VEP Migraineurs versus Controls. *Note*. No effects were significant at *p* < .05. Blank cells represent missing data.

Subgroup analysis comparing VEP P100 and N135 amplitudes for diagnosis of migraine without aura (MO) versus HC, migraine with aura (MA) versus HC, and MO versus MA showed that the P100 amplitude was significantly reduced for MO, with a small effect size across 11 studies (*g* = −0.26, 95% C.I. [−0.50, −0.03], *p* = .03, *I*^2^ = 34.83%, *p* = .12), see Figure 4, and MA (*g* = −0.30, 95% C.I. [−0.59, 0.00], *p* = .049, *I*^2^ = 26.25%, *p* = .24), see Figure 5, when compared to HC. Amplitude of P100 was not found to differ between MO and MA across six studies, *g* = 0.22, 95% C.I. [−0.24, 0.67], *p* = .35, *I*^2^ = 62.86%, *p* = .02 (Supplementary Figure 1a). This suggests diagnosis of MO or MA may not influence P100 amplitude during the interictal period, although more studies are required due to high heterogeneity.

**Figure 4.**
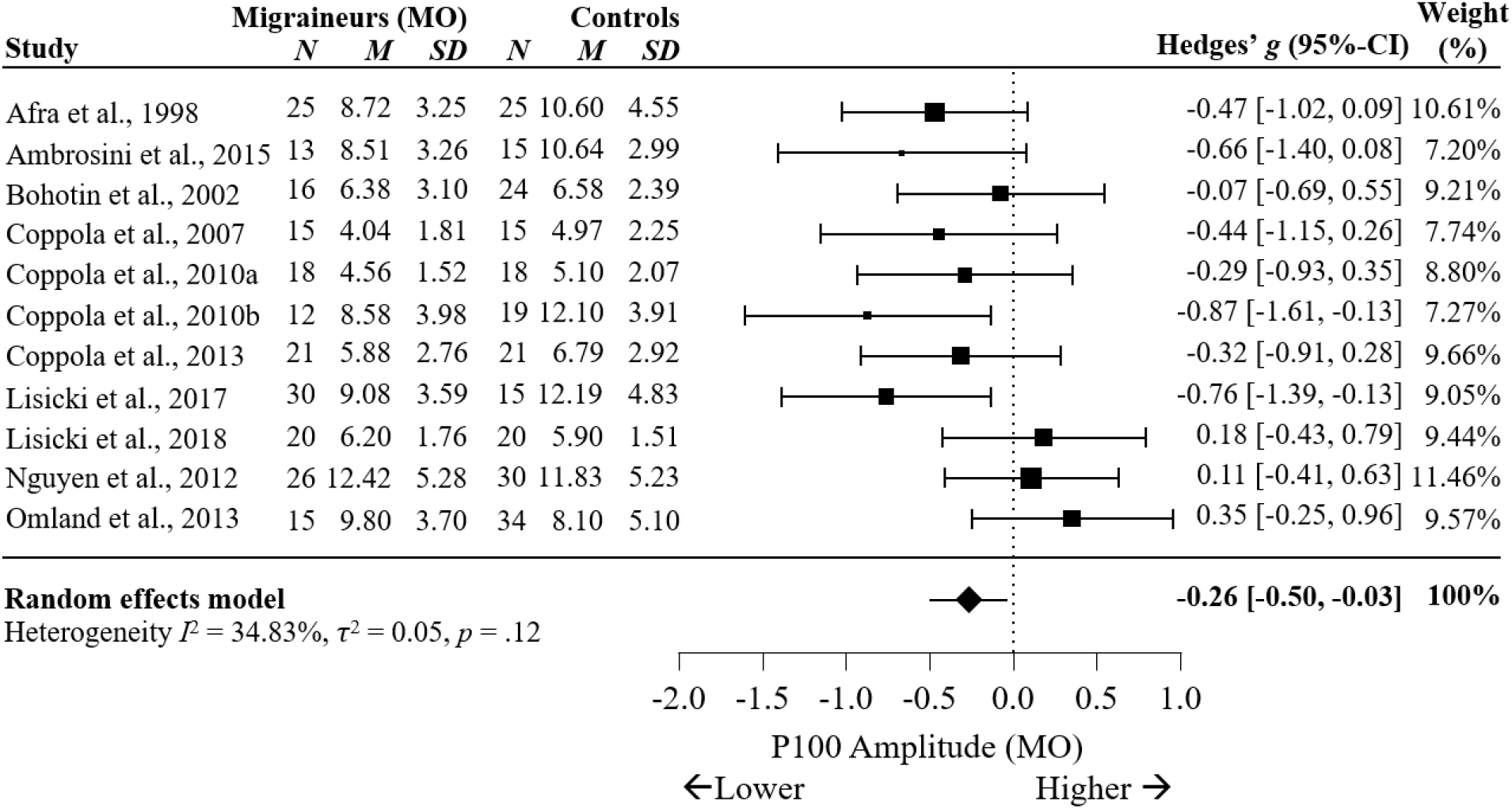
Forest Plots Comparing VEP P100 Amplitudes in Migraineurs Without Aura versus Controls

**Figure 5.**
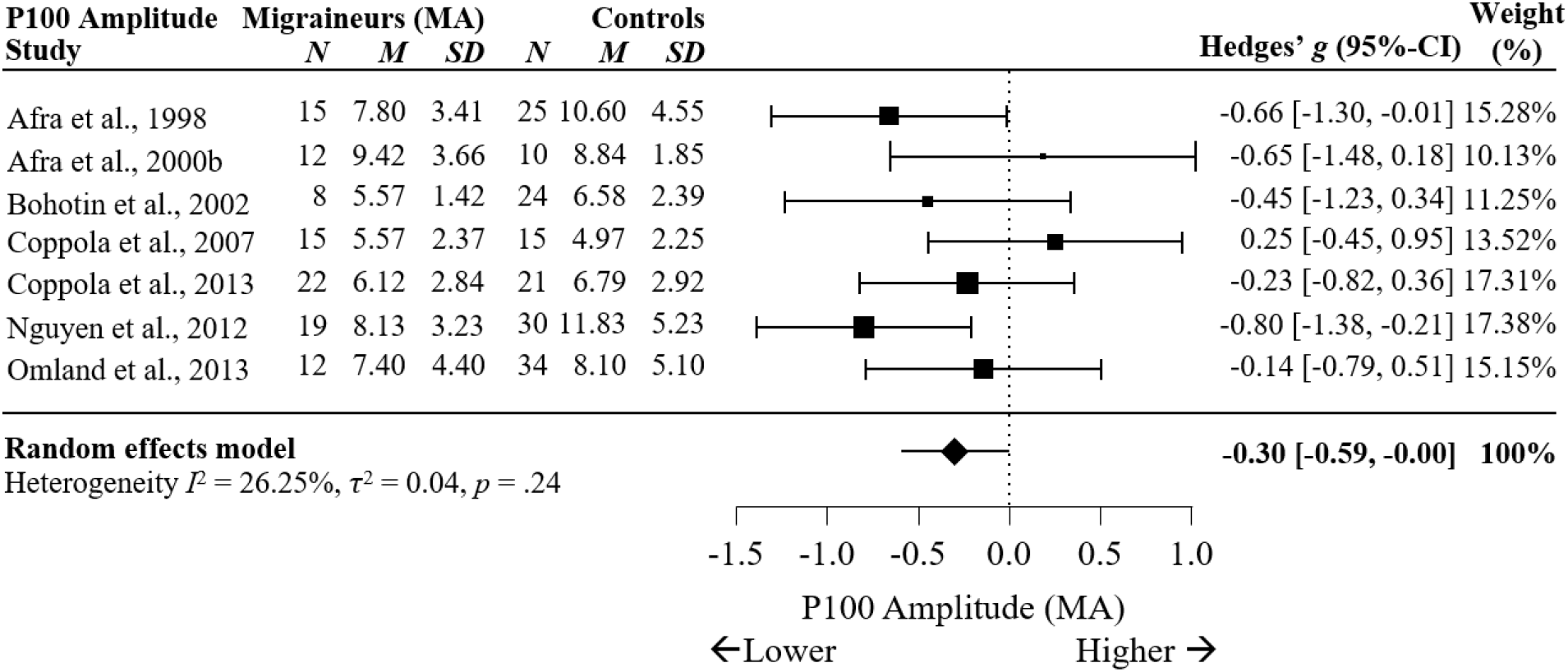
Forest Plots Comparing VEP P100 Amplitudes in Migraineurs With Aura versus Controls

No significant differences were seen for N135 amplitude between MO and HC across five studies, *g* = −0.29, 95% C.I. [−1.00, 0.42], *p* = .42, *I*^2^ = 84.86%, *p* < .001 (Supplementary Figure 2), MA and HC across four studies, *g* = −0.50, 95% C.I. [−1.16, 0.16], *p* = .14, *I*^2^ = 73.60%, *p* < .01 (Supplementary Figure 3), or MO and MA for N135 amplitude across four studies, *g* = 0.17, 95% C.I. [−0.17, 0.51], *p* = .34, *I*^2^ = 0.00%, *p* = .52 (Supplementary Figure 1b).

#### 3.4.2. Meta-Analyses for VEP Habituation in Migraine

Meta-analysis comparing differences between MG and HC in VEP P100 and N135 habituation showed significant differences in P100 habituation between MG and HC with a large effect size across 17 studies, *g* = 1.15, 95% C.I. [0.68, 1.62], *p* < .001, *I*^2^ = 93.61%, *p* < .001, see Figure 6. The positive meta-effect indicates migraineurs may have reduced habituation of P100 amplitude compared to non-migraineurs.

**Figure 6.**
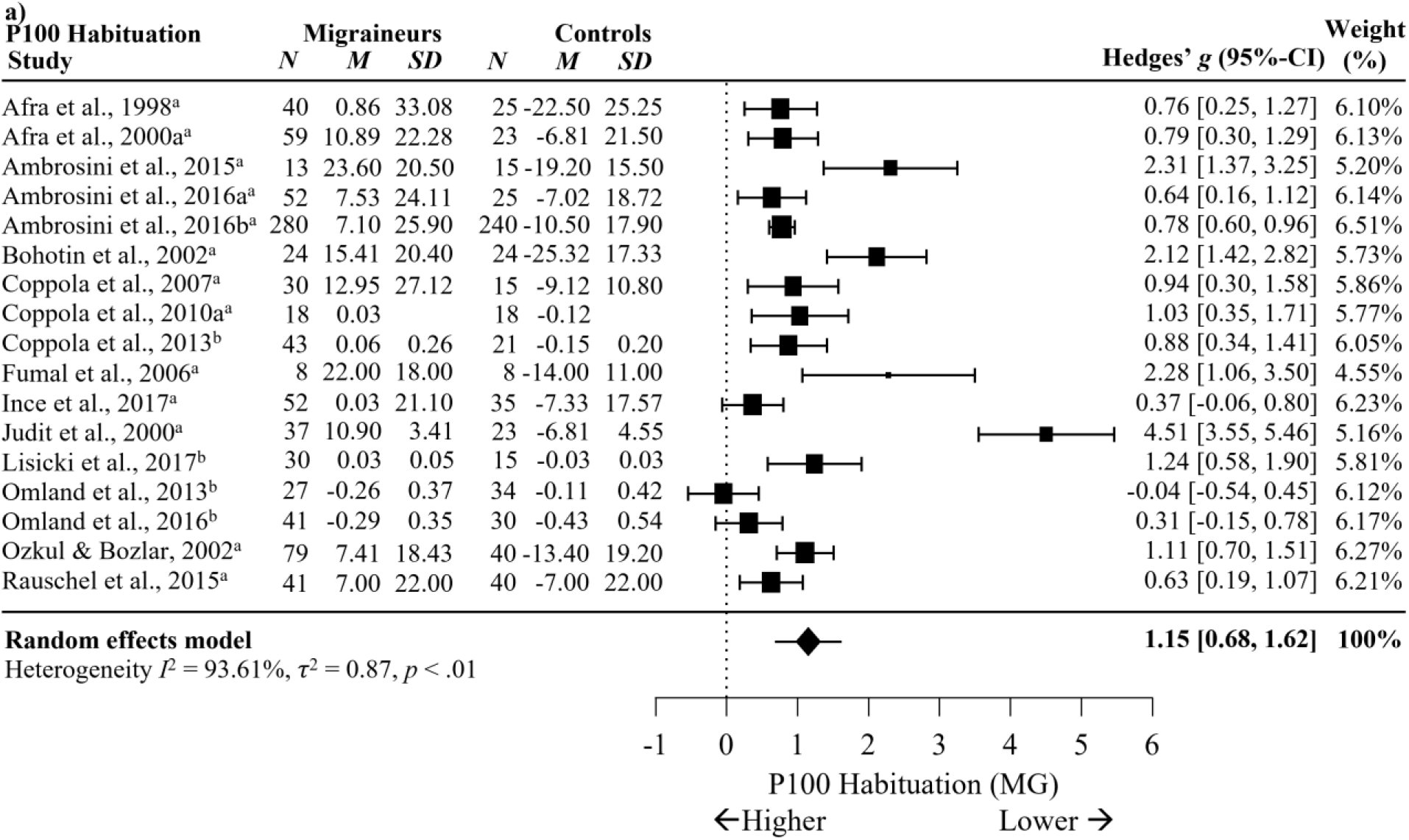
Forest Plots Comparing VEP P100 Habituation in Migraineurs versus Controls. *Note*. ^a^ = percentage difference, ^b^ = slope. Blank cells are missing data.

Subgroup analysis comparing MO and HC in VEP habituation for P100 showed significant differences between MO and HC in P100 habituation with a large effect size across 12 studies, *g* = 1.30, 95% C.I. [0.63, 1.97], *p* < .001, *I*^2^ = 92.42%, *p* < .001, see Figure 7. This suggests P100 habituation is largely reduced in migraineurs without aura compared to non-migraineurs, although high heterogeneity indicates moderating variables may impact this effect. Meta-regression showed check size significantly moderated the meta-effect calculation, *p* = .03, marginally reducing heterogeneity from *I*^2^ = 92.42% to *I*^2^ = 89.68%. This suggests variance in P100 habituation reported between studies is unlikely due to stimulus spatial frequency and may be due to other factors.

**Figure 7.**
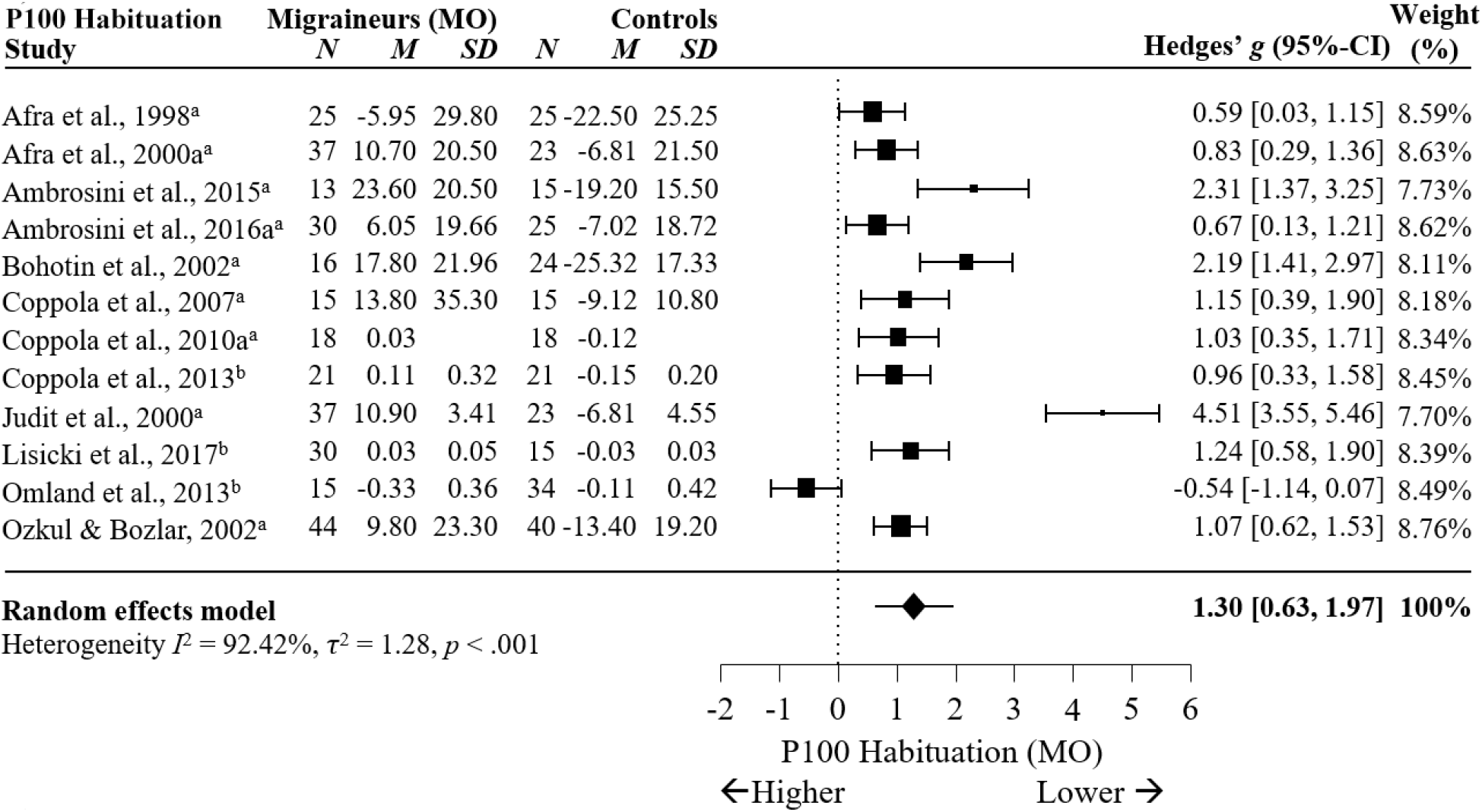
*Forest Plots Comparing VEP P100 Habituation* in *Migraineurs Without Aura versus Controls*. *Note*. ^a^ = percentage difference, ^b^ = slope. Blank cells are missing data.

Significant differences in P100 habituation between MA and HC with a large effect size across eight studies are shown in Figure 8, *g* = 0.88, 95% C.I. [0.52, 1.25], *p* < .001, *I*^2^ = 61.56%, *p* = .01. This suggests P100 habituation is also reduced in MA compared to controls, although high heterogeneity indicates moderating variables may impact this effect. Meta-regression showed check size significantly moderated the meta-effect calculation by reducing the influence of one outlier, *p* < .001, decreasing heterogeneity from *I*^2^ = 61.56% to *I*^2^ = 0.004%. Analysis of VEP habituation between migraineurs with and without aura showed no significant differences in P100 habituation between MO and MA across eight studies, *g* = 0.01, 95% C.I. [−0.22, 0.24], *p* = .92, *I*^2^ = 14.15%, *p* = .32 (Supplementary Figure 4a). This suggests migraineurs with and without aura consistently show no differences in P100 habituation.

**Figure 8.**
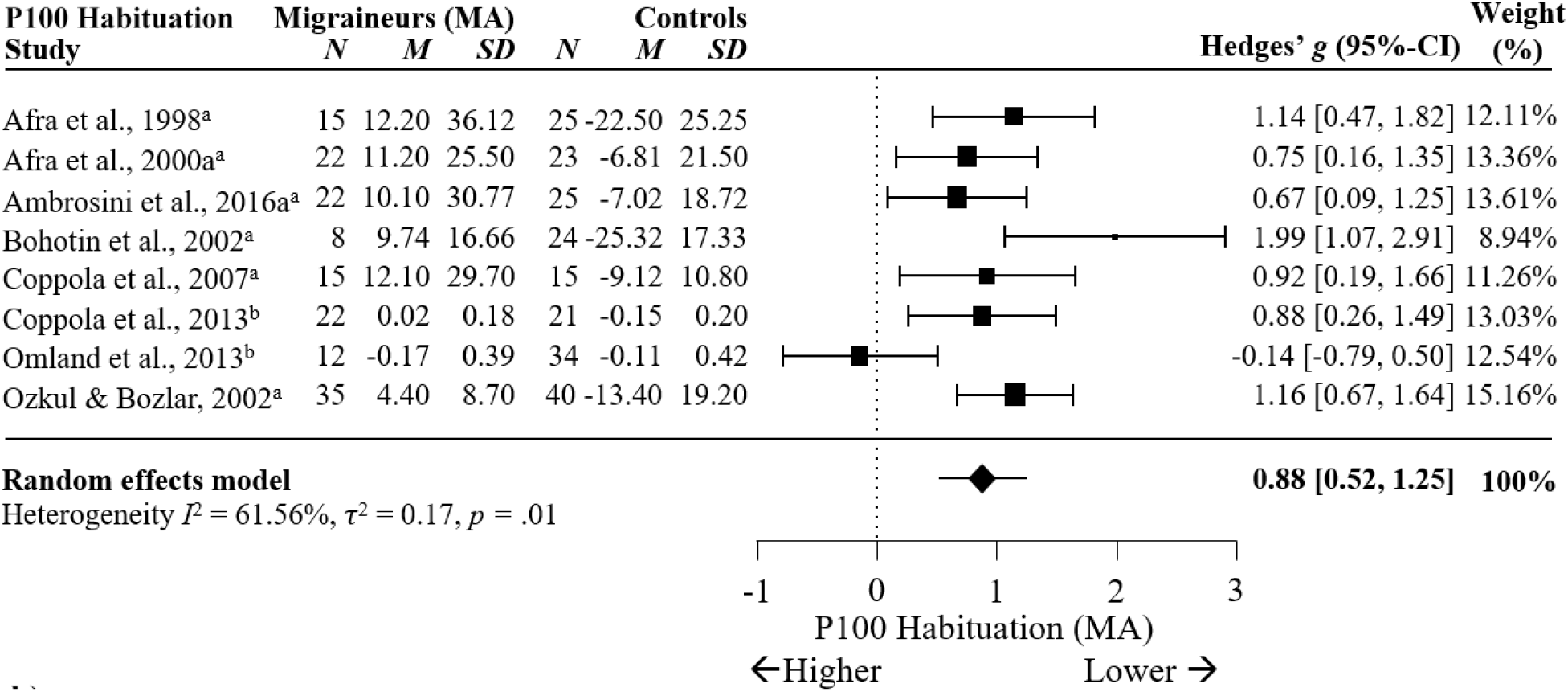
Forest Plots Comparing Migraineurs With Aura versus Controls in VEP P100 Habituation *Note*. ^a^ = percentage difference, ^b^ = slope.

Significant differences were also found in N135 habituation between MG and HC across five studies are shown in Figure 9, *g* = 1.09, 95% C.I. [0.13, 2.05], *p* = .03, *I*^2^ = 91.961%, *p* < .001. Although this suggests that migraineurs have largely reduced N135 habituation compared to non-migraineurs, this finding should be cautiously interpreted given the small number of studies analysed, wide confidence intervals and substantial heterogeneity.

**Figure 9.**
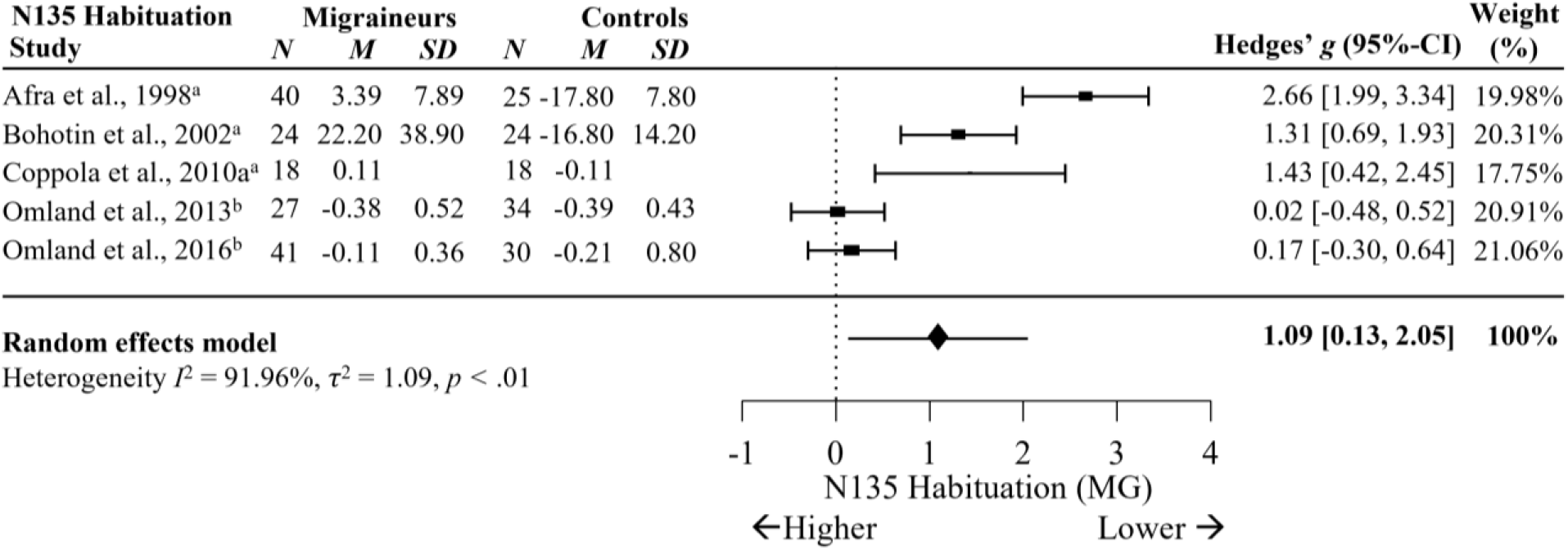
Forest Plots Comparing VEP N135 Habituation in Migraineurs versus Controls. *Note*. ^a^ = percentage difference; ^b^ = slope. Blank cells indicate missing data.

Significant differences in N135 habituation were observed between MO and HC with a large effect size across four studies, *g* = 1.59, 95% C.I. [0.54, 2.64], *p* < .01, *I*^2^ = 86.71%, *p* < .001, see Figure 10. Although this suggests N135 habituation appears largely reduced in migraineurs without aura compared to non-migraineurs, this finding should be cautiously interpreted given the small number of studies analysed, wide confidence intervals and substantial heterogeneity. There were no differences between MA and HC in N135 habituation across three studies, *g* = 1.02, 95% C.I. [−0.63, 2.67], *p* = .23, *I*^2^ = 92.87%, *p* < .001 (Supplementary Figure 5), nor for N135 habituation between MO and MA across three studies, *g* = 0.29, 95% C.I. [−0.47, 1.05], *p* = .45, *I*^2^ = 68.42%, *p* = .04 (Supplementary Figure 4b).

**Figure 10.**
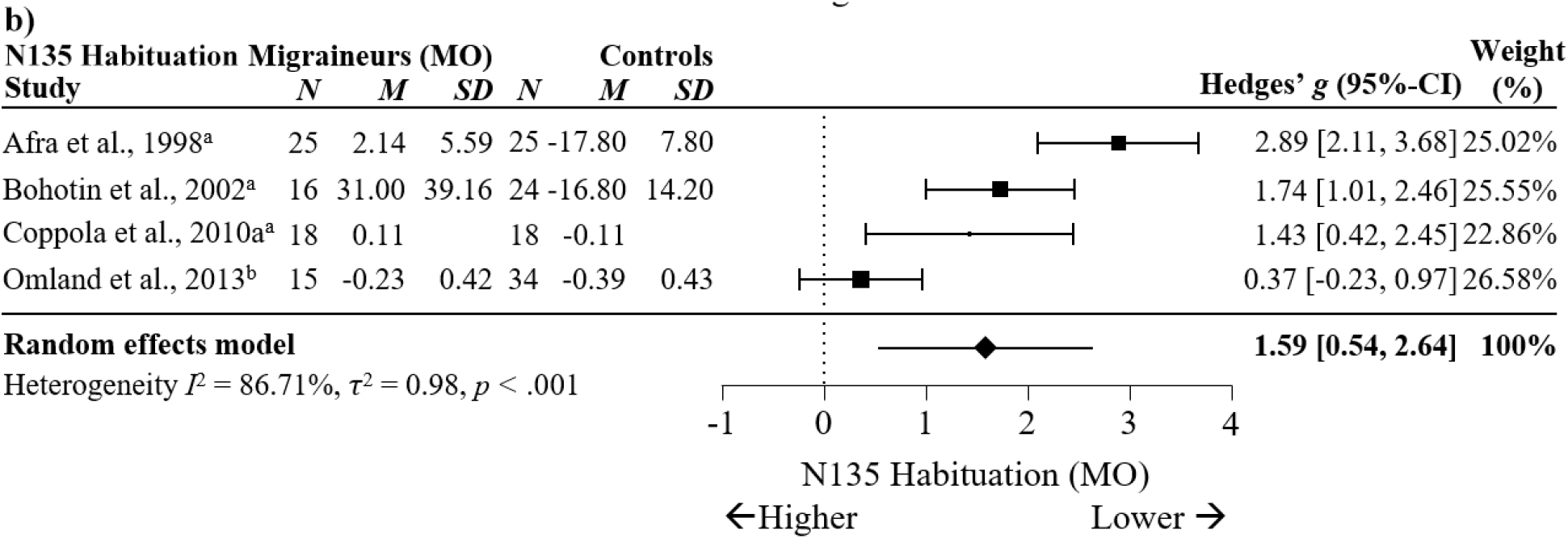
Forest Plots Comparing Migraineurs Without Aura versus Controls in VEP N135 Habituation *Note*. ^a^ = percentage difference, ^b^ = slope. Blank cells indicate missing data.

#### 3.5. Summary of I^2^ patterns and Funnel Plot Findings

Meta-analyses using studies that measured VEP P100 amplitudes tended to show moderate heterogeneity. By comparison, similar analyses of N135 amplitude showed substantial heterogeneity. Analyses of VEP P100 and N135 habituation often showed substantial heterogeneity. Of the analyses with sufficient number of studies included, assessments of publication bias showed mixed results. Refer to Supplementary Figures 6-13 for funnel plots and tests of asymmetry.

### 4. Discussion

This study aimed to meta-analyse case-control studies comparing visually evoked potentials (VEPs) between interictal migraineurs and non-migraine controls. Initial searches yielded 225 articles, of which 23 met the inclusion criteria and risk of bias assessment. The most important findings from the meta-analyses on the included studies was that migraineurs with and without aura showed slightly reduced P100 amplitude compared to non-migraine controls, but did not differ in N135 amplitudes. Furthermore, migraineurs showed largely reduced P100 and N135 habituation compared to controls. Only migraineurs without aura showed statistically reduced N135 habituation compared to controls, possibly because there were greater sample numbers for migraineurs without aura than migraineurs with aura. However, this result is confounded by the few statistically heterogeneous studies analysed. Finally, differences in VEP amplitudes and habituation were not found when comparing migraine subgroups of migraine with aura (MA) to migraine without aura (MO). Latency of VEPs were not analysed due to insufficient data available.

The primary hypothesis that migraineurs as a group would show altered VEP amplitudes during the interictal period compared to non-migraine controls was not supported by the 17 studies that met the criteria for a comparison of VEP amplitudes between all migraineurs and controls. However, in the 12 studies where sufficient subgroup data was present to allow meta-analysis, P100 amplitudes were reduced in the MA and MO subgroups when separately compared with controls. The second primary hypothesis was that migraineurs tested during the interictal period would show reduced VEP habituation to repeated stimulation compared to non-migraineurs, and this was partially supported with results demonstrating that all migraineurs (*N* = 17), including subgroups for MO (*N* = 12) and MA (*N* = 8), showed reduced P100 habituation compared to non-migraineurs. Although analysis of five studies found reduced N135 habituation in migraineurs compared to non-migraineurs, subgroup analysis found that only MO showed reduced N135 habituation compared to non-migraineurs in four studies.

The secondary aim of comparing visual function between migraine subgroups (MO compared with MA) demonstrated no differences in VEP amplitudes or habituation. Examination of the moderating effects of age, migraine frequency, years of migraine, and stimulus reversal rate or stimulus size on migraine VEPs was not possible due to insufficient number of studies providing data, lack of statistically matched samples and/or insufficient variability between studies.

#### 4.1. VEP Amplitudes during Interictal period of Migraine and Visual Processing

The present meta-analysis found reduced amplitude of visually evoked P100 waveform in the migraine MO and MA groups compared to non-migraineurs. It is important to note that (*i*) this finding does not appear to be robust against publication bias, with a small fail-safe *N* for both MO and MA groups suggesting that the significant findings could be negated by a few unpublished studies with non-significant results and (*ii*) a significant reduction in the P100 amplitude was not identified when comparing all migraineurs to controls (an analysis with larger sample size and greater statistical power). However, the subgroup analyses showed minimal heterogeneity, providing reasonable certainty that the findings were consistent across the literature.

These results suggest that migraineurs during the interictal period consistently experience a slight reduction in P100 amplitude, which is an early VEP component typically associated with the fast-conducting magnocellular pathway (M-pathway; Brown et al., 2018; Klistorner et al., 1997). Analysis of temporal processing, in particular VEP latency, was not possible as only four of the 23 studies examined this variable, despite the current results and earlier reports (O’Hare & Hibbard, 2016; Shepherd, 2019), suggesting that M-pathway processing in visual tasks requiring perception of object motion and orientation is impaired in migraineurs during the interictal period. Moreover, it is unclear whether reduced pattern-reversal VEPs are due to decreased excitatory mechanisms and/or increased inhibitory mechanisms (Cosentino et al., 2014), which remains an important gap in the literature (Vecchia & Pietrobon, 2012). Nevertheless, the finding of reduced VEP amplitude in the present meta-analysis may be interpreted as decreased M cell subcortical pathway recovery to repeated stimulation that contrasts with the prominent theory that migraine, particularly with aura, is characterised by increased cortical hyperexcitability to sensory stimuli during the interictal period (Barbanti et al., 2020).

The term “cortical excitability” is often used to describe electrophysiological differences observed in interictal migraineurs when compared with non-migraineurs (Aurora & Wilkinson, 2007; Chen et al., 2012). However, it is difficult to interpret how decreased VEP amplitudes observed in the present meta-analysis relates to physiological changes or an underlying cortical excitability in migraineurs. This is further complicated by differences in results across the different electrophysiology techniques used in migraine populations. For instance, our meta-analysis may be compared to the findings of a previous meta-analysis by Brigo et al. (2013) examining the effects of direct transcranial magnetic stimulation (TMS) on the primary visual cortex (V1), which found that migraineurs with aura, but not migraineurs without aura, had heightened excitability (as measured by increased phosphates) in V1 compared to non-migraineurs (Brigo et al., 2013). This may be explained by methodological differences between TMS and EEG, whereby TMS quantifies general cortical responsivity or arousal levels by directly stimulating V1 at variable stimulation levels (Magis et al., 2013), whereas VEPs instead quantify visual information processing between retina and V1 by recording electrical responses to visual stimulation from occipital electrodes (Odom et al., 2016). Thus, hyperexcitability defined by using direct TMS may reflect a lower threshold for visual cortex activation in migraine (Stankewitz & May, 2009), rather than increased responsivity to visual stimulation. In addition to these differences in how visual system activity is measured by TMS and VEP methodologies, the meta-analysis on TMS studies found substantial heterogeneity and did not restrict inclusion to studies recruiting interictal migraineurs (Brigo et al., 2013). As the present VEP meta-analysis found minimal heterogeneity and only included studies that explicitly included interictal migraineurs, our finding of reduced VEP amplitudes may better reflect M pathway neuronal recovery to repeated stimulation during visual processing in interictal migraineurs than TMS. Nevertheless, methodological limitations of standard checkerboard pattern-reversal VEP technology limit further interpretation of the findings of this meta-analysis until multifocal VEPS can be recorded and temporal analysis of conduction latencies can be measured (Klistorner et al., 1997).

Migraineurs have often been reported to experience visual discomfort or have migraines triggered when viewing certain visual stimuli during the interictal period (Peroutka, 2014; Shepherd, 2019), such as high-contrast black-and-white striped (Harle et al., 2006) and checkerboard patterns (Sand & Vingen, 2000). Defocusing during stimulation may also reduce VEP amplitudes (Nguyen et al., 2016), particularly for P100 (Creel, 2019). Thus, reduced VEP amplitudes in migraine could be due to migraineurs defocusing from discomfort during pattern-reversal stimulation (Nguyen et al., 2012). Although it is recommended that at least two blocks of VEP recordings are averaged to establish a reliable VEP response (Odom et al., 2016), most included studies only reported amplitudes from the first block of VEP recordings to minimise the influence of response habituation over prolonged stimulation (Magis et al., 2007). This raises concerns regarding the precision of results of the studies analysed that showed reduced VEP amplitudes in migraineurs. Nevertheless, our meta-analytic finding of reduced VEP amplitudes is consistent with previous reports of atypical habituation patterns in migraineurs (Magis et al., 2013). Ambrosini et al. (2003) theorised that low visual responsivity, as indicated by reduced VEP amplitudes or slower latency and slower neuronal recovery to repetitive stimulation in the first block of testing, could prevent migraineurs from reaching the threshold required to trigger normal habituation and subsequently a VEP amplitude decrement. This could be a protective mechanism preventing migraineurs from excessive neuronal excitation in response to visual stimulation, allowing greater cortical activation before the maximum threshold is reached (Nguyen et al., 2012). Thus, the reduced VEP amplitudes in migraineurs identified in the present meta-analysis aligns with the second major finding that VEP habituation was reduced in interictal migraineurs across the literature.

#### 4.2. VEP Habituation in Interictal Migraine – Impaired Filtering of Visual Information

The hypothesis that migraineurs possess atypical habituation patterns during the interictal period was supported by results, showing reduced habituation of P100 amplitude over time compared to non-migraineurs. Analysis of all migraineurs versus controls showed a large effect and substantial heterogeneity, while MO and MA subgroups showed large effects with substantial heterogeneity. Subgroups with MA showed minimal heterogeneity after meta-regression reduced the influence of one outlier. Together, results suggest that migraineurs during the interictal period may have a moderate to large impairment in neural habituation to repeated visual stimulation. Reduced P100 habituation and P100 amplitudes support a hypothesis that impaired visual processing in the fast-conducting M-pathway is potentially associated with migraine during the interictal period.

Habituation is considered a basic biological mechanism of learning and memory, as it enables cortical neurons to direct metabolic resources towards novel stimuli by filtering unimportant or familiar information (McDiarmid et al., 2017). Previous psychophysical studies have found that migraineurs perform poorly in tasks related to object motion and orientation that require filtering of visual noise during the interictal period (Shepherd, 2019; Tibber et al., 2014), as well as tasks that demand executive control of attention to visual stimuli (Han et al., 2019). These findings in conjunction with reduced VEP habituation of P100 amplitude may point to anomalies in M-pathway driven visual attention and processing in migraine during the interictal period.

Atypical habituation has also been associated with disruptions in cortical mechanisms responsible for decreasing excitation and/or increasing inhibition in response to visual stimulation (Ramaswami, 2014). Similar to reduced P100 amplitudes, it remains unclear how cortical excitation and inhibition mechanisms each contribute to reduced pattern-reversal VEP habituation in migraineurs during the interictal period (Cosentino et al., 2014). Furthermore, de Tommaso et al. (2014) have suggested that reduced habituation in migraineurs could instead reflect potentiation or increase in visual responses over time. Since included studies used between-groups designs and few/none include measures of anxiety (Al-Ezzi et al., 2020) or visual discomfort, it is not possible to precisely differentiate lack of VEP habituation from potentiation when using results from the current meta-analysis.

In contrast to heightened cortical responsivity to visual stimulation, some authors argue that reduced habituation observed in migraine could result from increased stimulus-induced electrical noise (O’Hare & Hibbard, 2016). It is also possible that VEPs are impacted by chronic anxiety often associated with migraine (Al-Ezzi et al., 2020). This may explain why other sensory recordings, such as auditory, somatosensory, and nociceptive evoked potentials, have also shown atypical habituation in migraineurs (reviewed in Brighina et al., 2009; Magis et al., 2013). However, at this time atypical habituation of VEPs of the P100 is the only generalizable indicator of altered cortical processing associated with episodic migraines.

A recent review has noted that reduced cortical habituation occurs across many non-migraine disorders, including autism spectrum disorders, schizophrenia and Parkinson’s disease (McDiarmid et al., 2017) and fits with the hypothesis of Stankewitz and May (2009) who have argued that habituation deficits are a consequence of repeated exposure to pain activation and likely anxiety/stress caused by migraine symptoms, rather than a specific vulnerability factor underlying migraine pathophysiology. Consequently, atypical VEP habituation may worsen for migraineurs with more frequent migraines or more years suffering untreated migraines (Nguyen et al., 2012). Unfortunately, as alluded to earlier, the present meta-analysis could not explore such modifiers due to insufficient data regarding participant’s migraine frequency or years experiencing migraines being included in study demographics. In addition to missing data, methodological confounds related to VEP habituation were another important shortcoming of the studies included.

#### 4.3. Limitations of this Meta-Analysis & Future Research Directions

The generalisability of the results from this meta-analysis should be considered in the context of several methodological and data limitations. The meta-analysis suffered from limited data collected for small numbers of patients, heterogeneity of all comparisons, incomplete data on migraine severity, age at time of testing, age at disease onset, duration of disorder, visual discomfort and anxiety. Very few included studies reported the early N75 component, which is the earliest pattern-reversal VEP component depicting primarily M-pathway contribution to V1 (Klistorner et al., 1997). Furthermore, VEP stimulus check sizes varied across studies, further confounding the contribution of the M-pathway and P-pathway to P100 amplitudes and habituation. As such, it is not possible to precisely characterise reduced P100 amplitudes and habituation by M-pathway dysfunction in migraineurs during interictal periods. Rather than using alternating check sizes in pattern-reversal stimuli, future studies should incorporate a multifocal VEP stimulus and temporal analysis, which allows temporal dissociation of the M-pathway and P-pathway contributions to the cortical VEP (Klistorner et al., 1997). In addition, insufficient VEP latency data to date is another key limitation of this study and that of a previous review by Ambrosini et al. (2003) and hence warrants further research and analysis. Thus, it remains unclear whether migraineurs possess normal visual attention and processing speed functions, or indeed experience M-pathway impairment that has been hypothesised from behavioural measures (Shepherd, 2019).

The substantial heterogeneity between studies which assessed VEP habituation in all migraineurs and subgroups with MO remains a limitation to understanding of attentional processing in migraineurs. Few studies reported controlling for potential modifiers of VEP habituation such as sensory adaptation/fatigue (McDiarmid et al., 2017; Rankin et al., 2009) or anxiety (Al-Ezzi et al., 2020). Furthermore, the studies included in the present meta-analyses used inconsistent measures of habituation, measured as either the percentage change in VEP amplitudes between the first and last blocks of testing or the linear regression slope of VEP amplitudes across all blocks. These indexes of habituation often produce very distinct results even when used on the same datasets (Omland et al., 2011) and have led McDiarmid et al. (2017) to recommends authors include raw data as well as both VEP habituation indexes, as each method pertains some limitation, with inclusion of both providing more comprehensive data for future meta-analyses. Nevertheless, the present meta-analysis helped resolve previous controversies in the literature by highlighting that habituation of the P100 wave of pattern reversal VEPs is generally reduced in migraineurs during the interictal period.

Another limitation of this meta-analysis is that the contribution of migraine severity to VEP amplitudes and habituation for all included studies could not be tested. Aside from the aforementioned missing data for migraine frequency and years suffering migraines, conclusions can only be generalised to episodic migraine since no studies recruiting chronic migraineurs were included in this meta-analysis. Preliminary research has suggested that chronic migraineurs show similar VEPs to non-migraineurs when recorded using magnetoencephalography (Chen et al., 2011). A further EEG and VEP study (Viganò et al., 2018) that did not meet our age inclusion criteria has also found that electrophysiological responses were similar between chronic migraineurs during the interictal period and non-migraineurs. As such, further research is needed that measures VEPs in chronic migraineurs while excluding older adults due to age related declines in VEP latency (Brown et al., 2019).

Participant age was another limitation of the present meta-analysis. Aging is associated with both attenuation of VEPs (Brown et al., 2019) as well as changes in migraine symptoms (Antonaci et al., 2014; Wijeratne et al., 2019). For example, one large study of migraineurs aged 16 to 80 found symptoms such as photophobia were less frequent in older adults (Kelman, 2006), presumably due to age-related ocular degeneration impacting the quality of the visual signal (reviewed in Brown et al., 2019). Thus, it is highly likely that age is a confounder of the results of this meta-analysis.

Lastly, a potential problem in the included studies was the dearth of objective information relating to the stage in the interictal period and migraine cycle. Recent evidence demonstrates that the early stages of migraine before prodromal symptoms emerge can be identified by daily vision tests from home (McKendrick et al., 2018). This highlights the need for further objective research regarding visual function during the interictal period and the inclusion of such measures in migraine VEP research.

### 5. Conclusion & Future Directions

This meta-analysis summarised studies that met the inclusion criteria regarding comparison of recording visually evoked potentials (VEPs) between migraineurs during the interictal period and non-migraine controls. The results indicate that migraineurs with and without aura, compared to non-migraineurs, showed slightly reduced P100 amplitudes and largely reduced P100 habituation to repeated stimulation during the interictal period. This suggests possible dysfunction in the fast-conducting magnocellular visual pathways that affect attention, motion detection and rates of visual processing in migraineurs during the interictal period. Although the generalisability of results is limited by missing data, participant confounds and inconsistent methodologies, these are unlikely to diminish the utility of abnormal VEPs as a physiological biomarker of cortical activation in migraineurs. Results from this meta-analysis highlight the likelihood of early magnocellular visual processing abnormalities in migraine, providing a platform for future experimental studies to advance migraine research by using temporal non-linear VEP analysis techniques specialised for separation of the magnocellular and parvocellular contributions to geniculostriate cortical visual processing.

## Supporting information

Suppelmentary Figure 5

* References marked with an asterisk were included in the meta-analysis.

