## Supplementary Table 1 for "Visual Processing During the Interictal Period Between Migraines: A Meta-Analysis"

*National Heart, Lung and Blood Institute Quality Assessment of Case-Control Studies (NHLBI, 2014)*

| **Criteria** | **Yes** | **No** | **Other (CD, NR, NA)** |
| --- | --- | --- | --- |
| 1. Was the research question or objective in this paper clearly stated and appropriate? |  |  |  |
| 2. Was the study population clearly specified and defined? |  |  |  |
| 3. Did the authors include a sample size justification? |  |  |  |
| 4. Were controls selected or recruited from the same or similar population that gave rise to the cases (including the same timeframe)? |  |  |  |
| 5. Were the definitions, inclusion and exclusion criteria, algorithms or processes used to identify or select cases and controls valid, reliable, and implemented consistently across all study participants? |  |  |  |
| 6. Were the cases clearly defined and differentiated from controls? |  |  |  |
| 7. If less than 100 percent of eligible cases and/or controls were selected for the study, were the cases and/or controls randomly selected from those eligible? |  |  |  |
| 8. Was there use of concurrent controls? |  |  |  |
| 9. Were the investigators able to confirm that the exposure/risk occurred prior to the development of the condition or event that defined a participant as a case? |  |  |  |
| 10. Were the measures of exposure/risk clearly defined, valid, reliable, and implemented consistently (including the same time period) across all study participants? |  |  |  |
| 11. Were the assessors of exposure/risk blinded to the case or control status of participants? |  |  |  |
| 12. Were key potential confounding variables measured and adjusted statistically in the analyses? If matching was used, did the investigators account for matching during study analysis? |  |  |  |
| Quality Rating (Good, Fair, or Poor) |  |  |  |
| Rater #1 Initials: |  |  |  |
| Rater #2 Initials: |  |  |  |
| Additional comments (If Poor, please state why): |  |  |  |

*Note.* CD = cannot determine, NR = not reported, NA = not applicable. Adapted from “Quality assessment of case-control studies” by National Heart, Lung, and Blood Institute, 2014, National Institutes of Health, (<https://www.nhlbi.nih.gov/health-topics/study-quality-assessment-tools>). Copyright 2014 by National Institutes of Health.

This tool assesses the following areas of bias: 1) Clear and appropriate research question; 2) Clearly defined study population; 3) Sample size justification; 4) Similarity/comparability of case and control samples; 5) Validity and reliability of case and control identification; 6) Cases clearly differentiated from controls; 7) Adequacy of random selection of case and control ; 8) Used concurrent controls; 9) Confirmation of the risk occurring prior to the development of the condition defining cases; 10) Clearly defined, valid, reliable and consistent definition of measures of exposure/risk; 11) Experimenters blinded to case/control status; 12) Analysis adjusted for confounding variables and accounted for matching. Two reviewers (TS and VN) independently assessed risk of bias for each study with discrepancies resolved via third senior reviewer (SC) consultation. Criteria responses were “yes”, “no”, or other (“cannot determine”, “not applicable”, “not reported”). Final quality was rated as “Good”, “Fair”, or “Poor”. Studies containing “fatal flaws” were deemed serious risk of bias and hence excluded from meta-analyses.
