## Supplementary Table 2 for "Visual Processing During the Interictal Period Between Migraines: A Meta-Analysis"

*Results from Risk of Bias Assessment (NHLBI, 2014)*

| Study | 1 | 2 | 3 | 4 | 5 | 6 | 7 | 8 | 9 | 10 | 11 | 12 | Rating |
| --- | --- | --- | --- | --- | --- | --- | --- | --- | --- | --- | --- | --- | --- |
| Afra et al., 2000a |  |  |  |  |  |  |  |  |  |  |  |  | Good |
| Afra et al., 1998 |  |  |  |  |  |  |  |  |  |  |  |  | Fair |
| Afra et al., 2000b |  |  |  |  |  |  |  |  |  |  |  |  | Good |
| Ambrosini et al., 2016a |  |  |  |  |  |  |  |  |  |  |  |  | Fair |
| Ambrosini et al., 2015 |  |  |  |  |  |  |  |  |  |  |  |  | Good |
| Ambrosini et al., 2016b |  |  |  |  |  |  |  |  |  |  |  |  | Fair |
| Bohotin et al., 2002 |  |  |  |  |  |  |  |  |  |  |  |  | Good |
| Coppola et al., 2007 |  |  |  |  |  |  |  |  |  |  |  |  | Good |
| Coppola et al., 2010a |  |  |  |  |  |  |  |  |  |  |  |  | Good |
| Coppola et al., 2010b |  |  |  |  |  |  |  |  |  |  |  |  | Fair |
| Coppola et al., 2013 |  |  |  |  |  |  |  |  |  |  |  |  | Good |
| Fumal et al., 2006 |  |  |  |  |  |  |  |  |  |  |  |  | Good |
| Ince et al., 2017 |  |  |  |  |  |  |  |  |  |  |  |  | Fair |
| Judit et al., 2000 |  |  |  |  |  |  |  |  |  |  |  |  | Poor |
| Lisicki et al., 2017 |  |  |  |  |  |  |  |  |  |  |  |  | Good |
| Lisicki et al., 2018 |  |  |  |  |  |  |  |  |  |  |  |  | Good |
| Logi et al., 2001 |  |  |  |  |  |  |  |  |  |  |  |  | Poor |
| Nguyen et al., 2012 |  |  |  |  |  |  |  |  |  |  |  |  | Good |
| Nguyen et al., 2014 |  |  |  |  |  |  |  |  |  |  |  |  | Good |
| Omland et al., 2013 |  |  |  |  |  |  |  |  |  |  |  |  | Good |
| Omland et al., 2016 |  |  |  |  |  |  |  |  |  |  |  |  | Good |
| Ozkul & Bozlar, 2002 |  |  |  |  |  |  |  |  |  |  |  |  | Fair |
| Rauschel et al., 2015 |  |  |  |  |  |  |  |  |  |  |  |  | Good |
| Sand et al., 2000 |  |  |  |  |  |  |  |  |  |  |  |  | Excluded |
| *Note.*  = Yes, = No, = Cannot determine, = Not reported, = Not applicable. | | | | | | | | | | | | | |
