## Supplementary Table 4 for "Visual Processing During the Interictal Period Between Migraines: A Meta-Analysis"

**Supplementary Table 3**

*Characteristics of Included Studies*

| Study | Demographics | | Migraine sample characteristics | | | | Stimulus details | | | Risk |
| --- | --- | --- | --- | --- | --- | --- | --- | --- | --- | --- |
|  | Sample sizes: *N*  (Sex: M, F) | Age:  *M* (*SD*), range | Dx tool | Migraines per month:  *M* (*SD*) | Migraine years:  *M* (*SD*) | Interictal period | Eye | Check size | rps |  |
| Afra et al., 1998 | MG: 40 (9M, 31F)  MO: 25  MA: 15  HC: 25 | MG: 36.00  HC: 30.00 | ICHD-1 |  |  | 5 days after last attack | Monocular | 0.13° | 3.10 | Fair |
| Afra et al., 2000a | MG: 59 (13M, 46F)  MO: 37  MA: 22  HC: 23 (5M, 18F) | MG: 35.00 (11.00)  HC: 27.00 (7.00) | ICHD-1 | MO: 4.00 (3.00)  MA: 2.00 (0.50) | MO: 15.00 (11.00)  MA: 14.00 (8.00) | 72 hours before and after | Monocular | 0.13° | 3.10 | Good |
| Afra et al., 2000b | MA: 12 (6M, 6F)  HC: 10 (2M, 8F) | MA: 34.00 (16.00)  HC: 28.00 (6.00) | ICHD-1 |  |  | 72 hours before and after | Monocular | 0.13° | 3.10 | Good |
| Ambrosini et al., 2015 | MA: 13 (6M, 7F)  HC: 15 (7M, 8F) | MO: 32.80 (10.20), 18-55  HC: 30.00 (7.90), 21-44 | ICHD-3 beta | MA: 3.20 (1.50) | MA: 11.70 (6.80) | 72 hours before and after | Monocular | 0.13° | 3.10 | Good |
| Ambrosini et al., 2016a | MG: 52 (8M, 44F)  MO: 30 (5M, 25F)  MA: 22 (3M, 19F)  HC: 25 (11M, 14F) | MO: 29.50 (8.00)  MA: 29.50 (9.00)  HC: 27.50 (8.10) | Clinical sample |  |  | 72 hours before and after | Monocular | 0.25° | 3.10 | Fair |
| Ambrosini et al., 2016b | MG: 280 (81M, 199F)  MO: 203  MA: 77  HC: 240 (102M, 139F) | MG: 32.04 (11.34)  HC: 27.26 (8.71) | ICHD-1 &  ICHD-2 |  |  | 72 hours before and after | Monocular | 0.25° | 3.10 | Fair |
| Bohotin et al., 2002 | MG: 24  MO: 16  MA:8  HC: 24 (10M, 14F) | MG: 33.50 (10.80)  HC: 23.50 (2.50) | ICHD-1 | MG: 2.31 (2.14) | MG: 13.71 (11.29) | 72 hours before and after | Monocular (right eye) | 0.13° | 3.10 | Good |
| Coppola et al., 2007 | MG: 30 (9M, 21F)  MO: 15 (5M, 10F)  MA: 15 (4M, 11F)  HC: 15 (6M, 9F) | MO: 31.00 (10.00),  20-53  MA: 30.00 (10.00),  19-56  HC: 27.70 (8.00), 21-49 | ICHD-2 | MO: 1.70 (1.50)  MA: 1.50 (1.20) |  | 72 hours before and after | Monocular (right eye) | 0.25° | 3.10 | Good |
| Coppola et al., 2010a | MO: 18 (7M, 11F)  HC: 18 (6M, 12F) | MO: 30.50 (9.50), 20-53  HC: 27.10 (7.70), 20-49 | ICHD-2 | MO: 2.00 (1.40) | MO: 18.00 (3.10) | 72 hours before and after | Monocular (right eye) | 0.25° | 3.10 | Good |
| Coppola et al., 2010b (personal communication) | MO: 12 (4M, 8F)  HC: 19 (9M, 10F) | MO: 28.10 (5.80), 24-36  HC: 26.20 (3.50), 19-33 | ICHD-2 | MO: 1.90 (2.10) | MO: 14.00 (5.10) | 72 hours before and after | Monocular | 0.25° | 3.10 | Fair |
| Coppola et al., 2013 | MG: 43 (7M, 36F)  MO: 21 (4M, 17F)  MA: 22 (3M, 19F)  HC: 21 (5M, 16F) | MO: 27.30 (6.50), 20-49  MA: 30.80 (9.70), 18-52  HC: 28.10 (7.60), 20-57 | Clinical sample | MO: 1.80 (1.10)  MA: 2.10 (2.30) | MO: 13.80 (9.50)  MA: 15.10 (8.80) | 72 hours before and after | Monocular (right eye) | 0.25° | 3.10 | Good |
| Fumal et al., 2006 | MG: 8 (3M, 5F)  MO: 6  MA: 2  HC: 8 (5M, 3F) | MG: 23.30 (1.00)  HC: 23.20 (1.90) | Clinical sample | MG: 1.80 (0.40) | MG: 5.00 (3.20) | 72 hours before and after | Monocular (right eye) | 0.13° | 3.10 | Good |
| Ince et al., 2017 | MG: 52 (9M, 43F)  MO: 48  MA: 4  HC: 35 (8M, 27F) | MG: 35.60 (9.10), 18-60  HC: 34.20 (9.60), 18-60 | ICHD-2 | MG: 7.60 (4.30) | MG: 6.90 (5.80) | 72 hours before and after | Monocular (left and right eyes) |  | 3.10 | Fair |
| Judit et al., 2000 | MO: 37 (3M, 34F)  HC: 23 (3M, 20F) | MO: 39.00  HC: 25.00 | ICHD-1 | MO: 4.00 | MO: 15.00 | 72 hours before and after | Monocular | 0.13° | 3.10 | Poor |
| Lisicki et al., 2017 | MO: 30 (6M, 24F)  HC: 15 (6M, 9F) | MO: 26.90 (6.99)  HC: 25.27 (3.43) | ICHD-3 beta |  |  | 72 hours before and after | Monocular (right eye) | 0.23° | 3.10 | Good |
| Lisicki et al., 2018 | MO: 20 (4M, 16F)  HC: 20 (5M, 15F) | MO: 32.20 (12.80)  HC: 34.80 (11.30) | ICHD-3 beta | MO: 4.10 (2.60) |  | 72 hours before and after | Monocular (right eye) | 1.13° | 3.10 | Good |
| Logi et al., 2001 | MG: 59 (10M, 49F)  MO: 40  MA: 19  HC: 30 (8M, 22F) | MG: 33.70 (14.90)  HC: 38.20 (9.60) | ICHD-1 |  |  | Around 10 days before | Monocular (left and right eyes) | 0.24° | 2.00 | Poor |
| Nguyen et al., 2012 | MG: 45 (9M, 36F)  MO: 26 (4M, 22F)  MA: 19 (5M, 14F)  HC: 30 (9M, 21F) | MO: 28.00 (6.00), 20-41  MA: 33.00 (6.00), 19-43  HC: 26.00 (7.00), 19-46 | ICHD-2 |  | MO: 13.00 (7.00)  MA: 19.00 (9.00) | 7 days before, 72 hours after | Binocular | 0.80° | 2.00 | Good |
| Nguyen et al., 2014 | MG: 17 (2M, 15F)  MO: 11  MA: 6  HC: 26 (8M, 18F) | MG: 29.00 (7.00), 19-43  HC: 25.00 (6.00), 19-46 | ICHD-2 |  | MO: 13.00 (6.00)  MA: 19.00 (10.00) | 7 days before, 72 hours after | Monocular | 0.80° | 2.00 | Good |
| Omland et al., 2013 | MG: 27 (3M, 24F)  MO: 15  MA: 12  HC: 30 (5M, 29F) | MG: 27.60 (8.40)  HC: 30.90 (10.40) | ICHD-2 | MG: 1.50 (0.60) | MG: 13.30 (7.60) | 48 hours before and after | Monocular (right eye) | 0.13°  &  1.08° | 3.00 | Good |
| Omland et al., 2016 | MG: 41 (5M, 36F)  MO: 24  MA: 2  MO&MA: 15  HC: 30 (5M, 25F) | MG: 38.50 (9.60), 19-56  HC: 37.80 (11.20), 21-59 | ICHD-2 | MG: 1.80 (0.60) | MG: 20.10 (9.60) | 48 hours before and after | Monocular (right eye) | 0.27° | 3.00 | Good |
| Ozkul & Bozlar, 2002 | MG: 79 (14M, 65F)  MO: 44  MA: 35  HC: 40 (11M, 29F) | MO: 36.00 (10.00)  MA: 34.00 (9.00)  HC: 33.00 (8.00) | ICHD-1 | MO: 3.50 (1.70)  MA: 3.10 (1.50) | MO: 13.00 (9.00)  MA: 12.00 (7.00) | 72 hours before and after | Monocular | 0.13° | 3.10 | Fair |
| Rauschel et al., 2015 | MG: 41 (3M, 38F)  MO: 38  MA: 3  HC: 40 (5M, 22F) | MG: 30.00 (10.00)  HC: 28.00 (8.00) | ICHD-3 beta | MG: 4.40 (2.30) | MG: 13.60 (9.90) | 48 hours before and after | Monocular (left eye) | 0.85° | 3.00 | Good |

*Note*. Sex: M, F = number of males and females, Dx tool = migraine diagnostic tool, Migraine years = years diagnosed with migraine, Eye = which eye(s) were stimulated during testing, Check size ° = stimulus size in minutes of arc, rps = reversals per second, Risk = risk of bias rating, MG = all migraineurs, MO = migraineurs with aura, MA = migraineurs with aura, HC = healthy controls, MO&MA: individuals diagnosed with both MO and MA, ICHD = International Classification of Headache Disorders. Blank cells represent missing data. Partially complete cells are indicative of missing data. Some studies only supplied demographics for MG, but not MO and MA individually and some studies supplied demographics for MO and MA individually but not combined.
