## Supplementary Table 5 for "Visual Processing During the Interictal Period Between Migraines: A Meta-Analysis"

**Supplementary Table 4**

*Individual VEP Results of Included Studies*

| Study | Stimulus details | | Amplitude | | Habituation | |
| --- | --- | --- | --- | --- | --- | --- |
|  | Blocks x Trials | VEP Component | Trials Averaged | Migraine and control group differences | Measures | Migraine and control group differences |
| Afra et al., 1998 | 15 x 100 | P100 &  N135 | First block | No significant differences between MO, MA and HC. | % Change | MO and MA had significantly reduced habituation than HC. |
| Afra et al., 2000a | 5 x 50 | P100 | First block | No significant differences between MO, MA and HC. | % Change | MO and MA had significantly reduced habituation than HC. |
| Afra et al., 2000b | 5 x 50 | P100 | All blocks | No significant differences between MA and HC. |  |  |
| Ambrosini et al., 2015 | 6 x 100 | P100 | First block | No significant differences between MO and HC. | % Change & Slope | MO had significantly reduced habituation than HC. |
| Ambrosini et al., 2016a | 6 x 100 | P100 |  |  | % Change | MG had significantly reduced habituation than HC. |
| Ambrosini et al., 2016b | 6 x 100 | P100 |  |  | % Change | MO and MA had significantly reduced habituation than HC. |
| Bohotin et al., 2002 | 6 x 100 | P100 &  N135 | First Block | Not reported. | % Change  & Slope | HC showed significant habituation, but MO and MA did not. |
| Coppola et al., 2007 | 6 x 100 | P100 | First block | No significant differences between MO, MA and HC. | % Change | MO and MA had significantly reduced habituation than HC. |
| Coppola et al., 2010a | 6 x 100 | P100 &  N135 | First Block | No significant differences between MO and HC. | % Change  & Slope | MO had significantly reduced habituation than HC. |
| Coppola et al., 2010b | 6 x 100 | P100 | First Block | Not reported. | Slope | Not reported. |
| Coppola et al., 2013 | 6 x 100 | P100 &  N135 | First Block | No significant differences between MO, MA and HC. | Slope | MO and MA had significantly reduced habituation than HC. |
| Fumal et al., 2006 | 6 x 100 | P100 |  |  | % Change | Not reported. |
| Ince et al., 2017 | 10 x 100 | P100 &  N135 | Sum of blocks 1, 5 & 10 | Not reported. | % Change | Not reported. |
| Judit et al., 2000 | 5 x 50 | P100 | Not reported | No significant differences between MO and HC. | % Change | Not reported . |
| Lisicki et al., 2017 | 6 x 100 | P100 | First block | MO had significantly reduced amplitude than HC. | Slope | MO had significantly reduced habituation than HC . |
| Lisicki et al., 2018 | 1 x 600 | P100 | First block | No significant differences between MG and HC. |  |  |
| Logi et al., 2001 | 2-3 x 100 | P100 | Unclear | No significant differences between MG and HC. |  |  |
| Nguyen et al., 2012 | 2 x 100 | P100 &  N135 | All blocks | MA had significantly reduced amplitude than HC. MO did not differ to HC. |  |  |
| Nguyen et al., 2014 | 1 x 200 | P100 | All blocks | No significant differences between MG and HC. |  |  |
| Omland et al., 2013 | 6 x 100 | P100 &  N135 | All blocks | MO had higher N135 amplitude than MA. MO and MA did not differ to HC. | % Change & Slope | No significant differences between MO, MA and HC. |
| Omland et al., 2016 | 6 x 100 | P100 &  N135 | All blocks | No significant differences between MO, MA and HC. | % Change & Slope | No significant differences between MO, MA and HC. |
| Ozkul & Bozlar, 2002 | 5 x 50 | P100 | First block | No significant differences between MO, MA and HC. | % Change | MO and MA had significantly reduced habituation than HC. |
| Rauschel et al., 2015 | 6 x 75 | P100 | First block | No significant differences between MG and HC. | % Change & Slope | MG had significantly reduced habituation than HC measured using slope, but not % change. |

*Note*. MG = all migraineurs, MO = migraineurs without aura, MA = migraineurs with aura, HC = healthy controls, % change = percentage change in VEP amplitude between first and last block of testing, slope = regression slope of the change in VEP amplitude across all blocks of testing. Blank cells represent missing data.
